# Glycolipid composition of the heterocyst envelope of *Anabaena* sp. PCC 7120 is crucial for diazotrophic growth and relies on the UDP-galactose 4-epimerase HgdA

**DOI:** 10.1101/489914

**Authors:** Dmitry Shvarev, Carolina N. Nishi, Iris Maldener

**Author notes:** corresponding author: Iris Maldener, Organismic Interactions, Interfaculty Institute of Microbiology and Infection Medicine, Eberhard Karls University of Tübingen, Auf der Morgenstelle 28, 72076 Tübingen, Germany.

## Abstract

The nitrogenase complex in the heterocysts of the filamentous freshwater cyanobacterium *Anabaena* sp. PCC 7120 fixes atmospheric nitrogen to allow diazotrophic growth. The heterocyst cell envelope protects the nitrogenase from oxygen and consists of a polysaccharide and a glycolipid layer that are formed by a complex process involving the recruitment of different proteins. Here we studied the function of the putative nucleoside-diphosphate-sugar epimerase HgdA, which along with HgdB and HgdC is essential for deposition of the glycolipid layer and growth without a combined nitrogen source. Using site-directed mutagenesis and single homologous recombination approach, we performed a thoroughly functional characterization of HgdA and confirmed that the glycolipid layer of the *hgdA* mutant heterocyst is aberrant as shown by transmission electron microscopy and chemical analysis. The *hgdA* gene was expressed during late stages of the heterocyst differentiation. GFP-tagged HgdA protein localized inside the heterocysts. The purified HgdA protein had UDP-galactose 4-epimerase activity *in vitro*. This enzyme could be responsible for synthesis of heterocyst-specific glycolipid precursors, which could be transported over the cell wall by the ABC transporter components HgdB/HgdC.

**Importance:** The phototrophic multicellular bacterium *Anabaena* sp. PCC 7120 is an important model organism for investigations of critical processes such as photosynthesis, nitrogen fixation, and cell differentiation in prokaryotes. The cyanobacterium utilizes specific cells, called heterocysts, to fix atmospheric nitrogen. Heterocyst function requires a specific cell envelope. We focused on unstudied aspects of heterocyst envelope formation and revealed the role of the UDP-galactose 4-epimerase HgdA in this process.

## Introduction

*Anabaena* sp. PCC 7120 (also known as *Nostoc* sp. PCC 7120, hereafter *Anabaena* sp.) belongs to a group of multicellular filamentous cyanobacteria that can differentiate and form heterocysts, cells specialized in N_2_ fixation. Upon removal of a source of combined nitrogen, heterocysts arise along the filaments in a semi-regular pattern, with approximately one heterocyst to ten vegetative cells.

Heterocysts host the extremely oxygen-sensitive nitrogenase complex (1–5). The required microoxic environment in the differentiating cells is achieved by shutting down of oxygenic photosynthesis, activation of respiration, and several morphological changes. The most obvious cellular modification is the synthesis of the heterocyst cell envelope outside of the normal Gram-negative cell wall (1–4). This heterocyst envelope consists of two different layers: the outermost exopolysaccharide (hep) layer and the underlying glycolipid (hgl) layer. The hgl layer restricts gas influx into the heterocyst cytoplasm, and the hep layer mechanically supports the hgl layer (2). The glycolipids of the hgl layer (HGLs) are heterocyst-specific and can differ in the aglycone length, sugar moiety, or number and type of functional groups (e.g., diol, keto-ol, and triol) (6–13).

In *Anabaena* sp., the most abundant HGLs are 1-(O-α-D-glucopyranosyl)-3,25-hexacosanediol (HGL_26_ diol) and its 3-ketotautomer (HGL_26_ keto-ol) (8, 13). The synthesis of HGLs and deposition of the hgl layer probably constitute a multi-step pathway involving products of different genes (2, 14), and many questions remain open. It is known that a type I secretion system (T1SS)-like transporter is involved in the efflux of HGLs from the inside of the developing heterocysts to form the hgl layer (15–18). This transporter is composed of the TolC homologue outer membrane protein HgdD, the periplasmic membrane fusion protein DevB, and the inner membrane ABC transporter DevCA. The DevBCA-HgdD efflux pump is essential for the hgl layer formation and heterocyst function (16, 17).

Several homologs of the *devBCA* gene cluster in the genome of *Anabaena* sp. have been identified (19, 20). Some are important for diazotrophic growth and heterocyst maturation (21– 23). The cluster *all5347/all5346/all5345* (*hgdB/hgdC/hgdA*) is of particular interest because the ATPase-coding gene *devA* is replaced by the *hgdA* gene coding for a putative epimerase. This gene cluster is essential for proper hgl layer deposition and growth of *Anabaena* sp. without combined nitrogen source (22, 23), but the functions of the protein HgdA (All5345) remain unknown.

Epimerases form a large group of enzymes that can be found in bacteria, animals and plants (24). They take part in important metabolic processes, for example, UDP-galactose 4-epimerase participates in the Leloir pathway, in which it converts UDP-galactose to UDP-glucose (25–27). Epimerases mainly constitute dimers, however other oligomeric states can also be found; the structures of some of these proteins have been resolved (28–31). In the present study, we investigated the role of the putative epimerase HgdA in *Anabaena* sp. during diazotrophic growth.

## Results

### The *hgdA* gene product is homologous to NDP-sugar epimerases

The *hgdA* gene is the third gene in the previously described cluster involved in heterocyst formation, *all5347/all5346/all5345* (*hgdB/hgdC/hgdA*) [Fig. 1A; (22, 23)]. It encodes a protein of 333 amino acids with a predicted molecular mass of 36.7 kDa. According to the results of a search using the NCBI BLAST tool, HgdA is a putative nucleoside-diphosphate-sugar epimerase belonging to the NAD-dependent epimerase/dehydratase superfamily and to the short-chain dehydrogenases/reductases (SDR) superfamily. An additional *in silico* search for homologs of HgdA using the PaperBLAST tool, which searches for homologs of a given protein in published articles (32), revealed SDRs and epimerases similar to HgdA (Fig. S1). However, according to the phylogenetic tree built by the web service Phylogeny.fr (33, 34), based on multiple sequence alignments of selected HgdA homologs found by PaperBLAST, HgdA is more closely related to epimerases (Fig. 1B).

**Figure 1.**
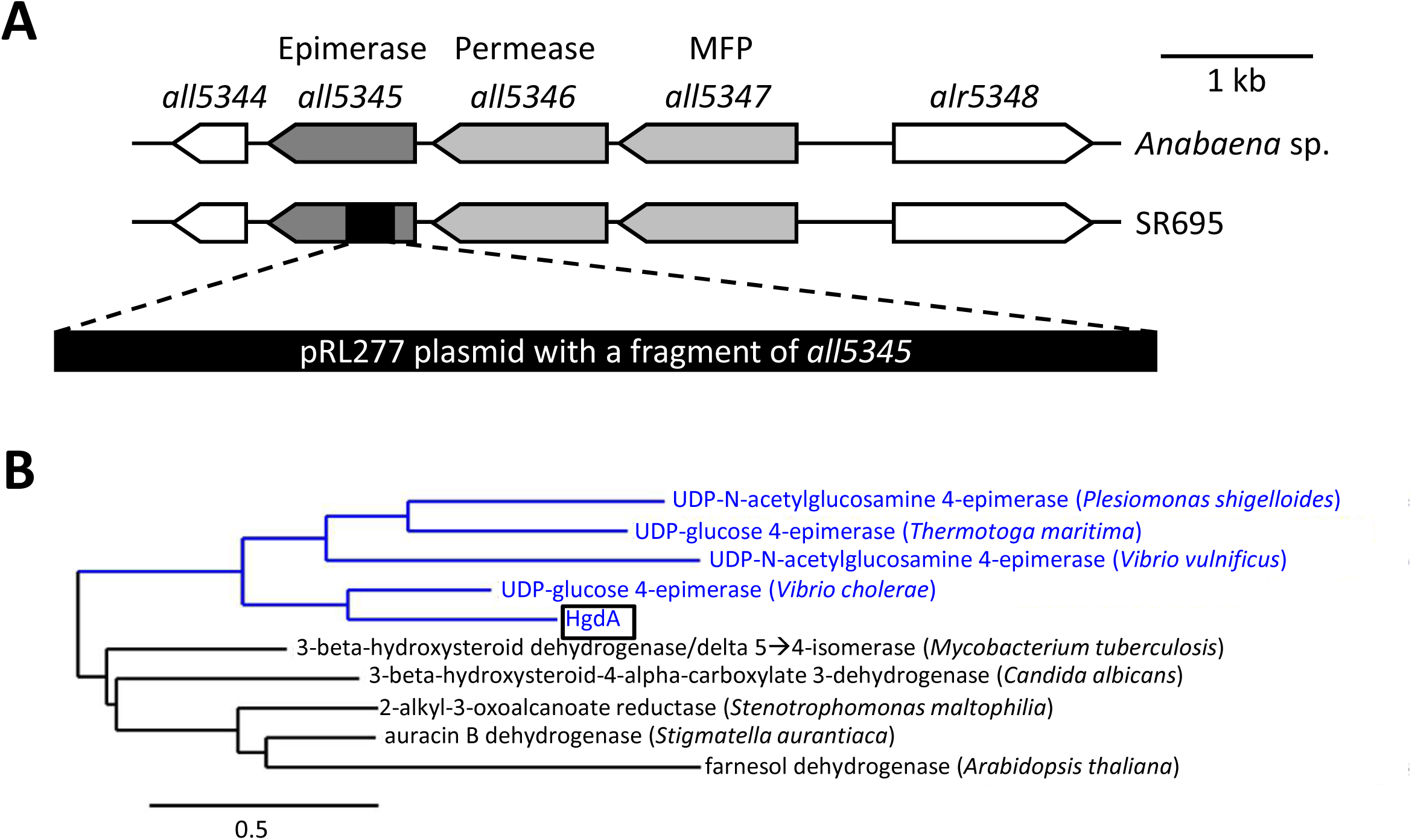
*in silico* analysis of HgdA. **A)** Genomic organization of the *Anabaena* sp. wild-type and the SR695 mutant strain in the region of the *hgdB/hgdC/hgdA* (*all5347/all5346/all5345*) gene cluster. MFP, membrane fusion protein. **B)** Phylogenetic tree of several characterized HgdA homologs found using the PaperBLAST online tool. The maximum-likelihood tree was constructed with the PhyML program (v3.1/3.0 aLRT) by the web service Phylogeny.fr.

### HgdA protein localizes specifically to heterocysts

To study the function of HgdA, we used semi-quantitative RT-PCR to follow the expression of the *hgdA* gene at different time points after transfer of a culture grown on NH_4_^+^ to medium without a combined nitrogen source (nitrogen stepdown) to induce heterocyst differentiation (Fig. 2A). The *hgdA* transcript levels were significantly higher at later stages of heterocyst formation, especially at 24–48 h after nitrogen stepdown. At these time points, the heterocysts were already visible by light microscopy. The up-regulation of *hgdA* was notably later than the previously reported up-regulation of *devB* (18), and the expression pattern of *hgdA* was almost identical to that of *hgdB* (23).

**Figure 2.**
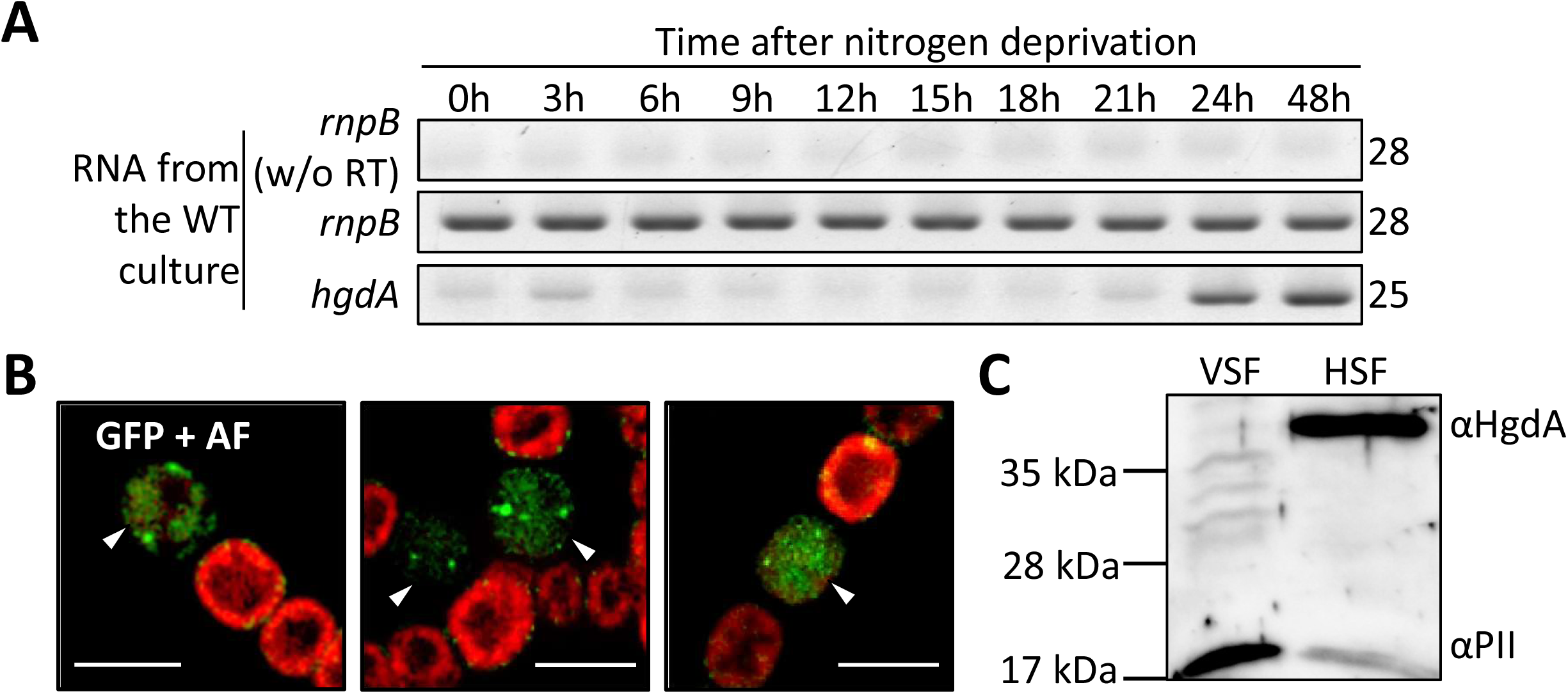
Analysis of *hgdA* expression in *Anabaena* sp. **A)** RT-PCR analysis of time-dependent *hgdA* expression. *rnpB,* ribonuclease B, used to ensure that the same amounts of RNA were used for cDNA synthesis in all samples. Numbers at the right indicate the number of PCR cycles. **B)** Localization of HgdA in cell compartments of *Anabaena* sp. Fluorescent micrographs of filaments bearing translational fusions of HgdA (All5345) with sfGFP after three days of nitrogen starvation. Green, GFP fluorescence; red, cyanobacterial autofluorescence (AF), white arrowheads, heterocysts. Bar, 5 µm. **C)** Western blot analysis of HgdA in vegetative cell fractions (VSF) and heterocyst soluble fractions (HSF). For comparison, antibodies raised against PII were used.

Since *hgdA* was only expressed under nitrogen starvation, we investigated the localization of the HgdA protein in diazotrophically grown filaments using a fusion protein consisting of HgdA linked at the C-terminus with sfGFP. The fusion protein localized solely to mature heterocysts. GFP fluorescence was equally distributed within the heterocyst and sometimes formed small foci (Fig. 2B). This observation is in line with the prediction that HgdA is an epimerase, which is a soluble enzyme.

In Western blot analysis, a protein cross-reacting with the HgdA-specific antibody was only visible in the sample obtained from the soluble heterocyst fraction, but not in the vegetative cell fraction (Fig. 2C) or in membrane fractions (not shown). The PII protein, used as a control, was detected in both samples, but was much more abundant in the vegetative cell fraction (Fig. 2C), which is in line with Paz-Yepez et al. (35), who demonstrated down-regulation of the PII-encoding gene *glnB* in *Anabaena* sp. heterocysts.

### The *hgdA* gene is essential for diazotrophic growth

To investigate the function of the *hgdA* gene in more detail, we created a mutant of this gene in *Anabaena* sp. by inserting an antibiotic resistance gene via homologous recombination (Fig. 1A, S2A). The mutant, SR695, was completely segregated (Fig. S2B). In medium with a combined nitrogen source, mutant SR695 did not differ from the wild-type in cell and filament morphology or in growth. However, the mutant was not able to grow diazotrophically, even though the filaments formed heterocysts after nitrogen stepdown (Fig. 3A, B, D). The mutant SR695 was complemented by introducing the self-replicating vector pIM612 (36) carrying the full-length *hgdA* sequence under control of the P_*glnA*_ promoter (37). The complemented mutant (SR695c) clearly grew better than the mutant SR695 at 7 days after nitrogen stepdown (Fig. 3A).

**Figure 3.**
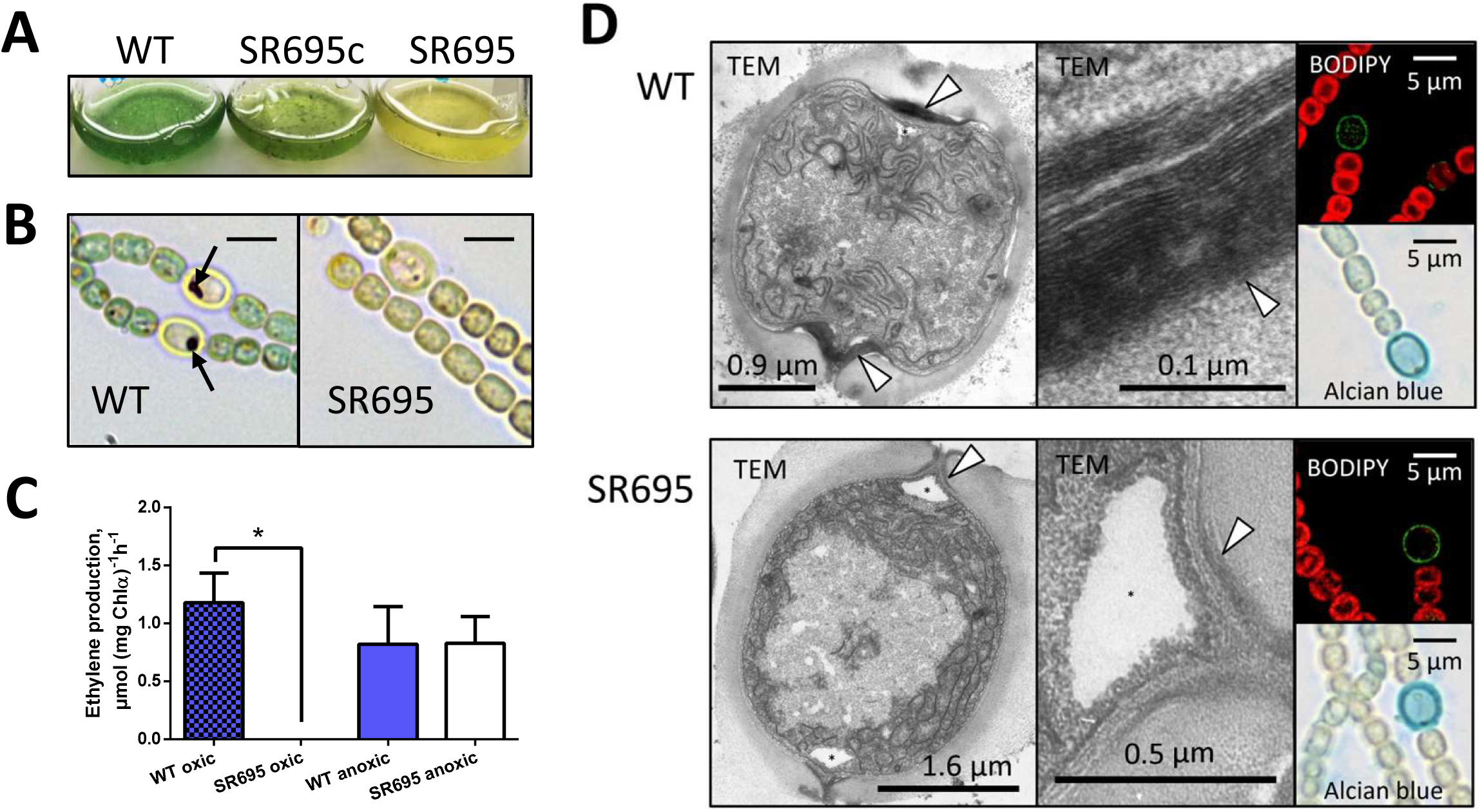
Phenotypic analysis of mutant SR695. **A)** Growth of wild-type (WT), mutant SR695, and the mutant complemented with the *hgdA* gene (SR695c) in liquid medium for 3 days without combined nitrogen source. **B)** Triphenyl tetrazolium chloride (TTC) staining of filaments of wild-type and mutant SR695. Arrows, dark formazan crystals of reduced TTC under microoxic and reducing conditions necessary for nitrogenase complex activity. Bars, 4.5 µm. **C)** Nitrogenase activity in wild-type and mutant SR695 cells under oxic and anoxic conditions assayed by acetylene reduction. Shown are mean values ± standard deviation of two experimental replicates. *, *P* < 0.05, Student’s *t*-test. **D)** Heterocyst cell envelope. Left panels, transmission electron micrographs (TEM) of wild-type and mutant SR695 heterocysts; white arrowhead, hgl layer; star, cyanophycin granule; right panels, micrographs of wild-type and mutant SR695 stained with BODIPY or alcian blue.

### The aberrant cell envelope of heterocysts of mutant SR695 cannot provide microoxic conditions for nitrogenase activity

We investigated whether heterocysts of the mutant SR695 provide microoxic conditions necessary for nitrogenase activity. We incubated mutant and wild-type cultures with triphenyl tetrazolium chloride (TTC) (38) and observed the dark crystals of reduced TTC only in wild-type heterocysts (Fig. 3B). Lack of dark TTC crystals in mutant heterocysts indicated that their inner environment was oxic.

Mutants with defects in heterocyst envelope layers only have nitrogenase activity when incubated under anoxic conditions (39). We assayed nitrogenase activity under oxic and anoxic conditions based on the measurement of acetylene reduction (40). The mutant SR695 had nitrogenase activity only under anoxic conditions, whereas the wild-type had nitrogenase activity under both oxic and anoxic conditions (Fig. 3C). Hence, the mutant heterocysts did not provide the microoxic conditions required for nitrogenase activity.

We analyzed the heterocyst envelope in more detail. We were able to detect both cell layers of the mutant heterocysts with standard labeling methods and light microscopy (41, 42), and the heterocysts appeared normal at this resolution (Fig. 3D). However, wild-type and mutant heterocysts differed in ultrathin sections analyzed by transmission electron microscopy (TEM). The mutant heterocysts lacked the laminated hgl layer, and the heterocyst–vegetative cell connections at the polar neck regions were thinner (Fig. 3D). These observations are comparable with those of the *hgdA* mutant FQ1647 (22). We did not find any structural differences in the hep layer or in the ultrastructure of vegetative cells between wild-type and mutant SR695.

We analyzed the glycolipid composition of mutant and wild-type hgl layers using thin-layer chromatography (TLC) of methanol extracts of both strains after nitrogen stepdown. We analyzed the content at two temperatures (20 and 28 °C) because the ratio of the major HGL forms can vary at different temperatures (43, 44). Both wild-type and mutant extracts contained the major HGLs, HGL_26_ keto-ol and HGL_26_ diol (45) at both temperatures (Fig. 4). However, the diol:keto-ol ratio of wild-type and mutant SR695 differed. At 28 °C, the diol:keto-ol ratio of wild-type heterocysts was higher than that of the mutant (Fig. 4). At 20 °C, the wild-type contained more of the keto-ol form than at 28 °C. Nevertheless, the wild-type was still different from the mutant, which did not show a temperature dependent ratio change (Fig. 4). As previously reported, a mutant in the upstream gene *hgdB* shows a similar phenotype at 28 °C (23). We also found that at 20 °C, differences in growth between wild-type and mutant SR695 were not as prominent as at 28 °C (Fig. 4, lower panels).

**Figure 4.**
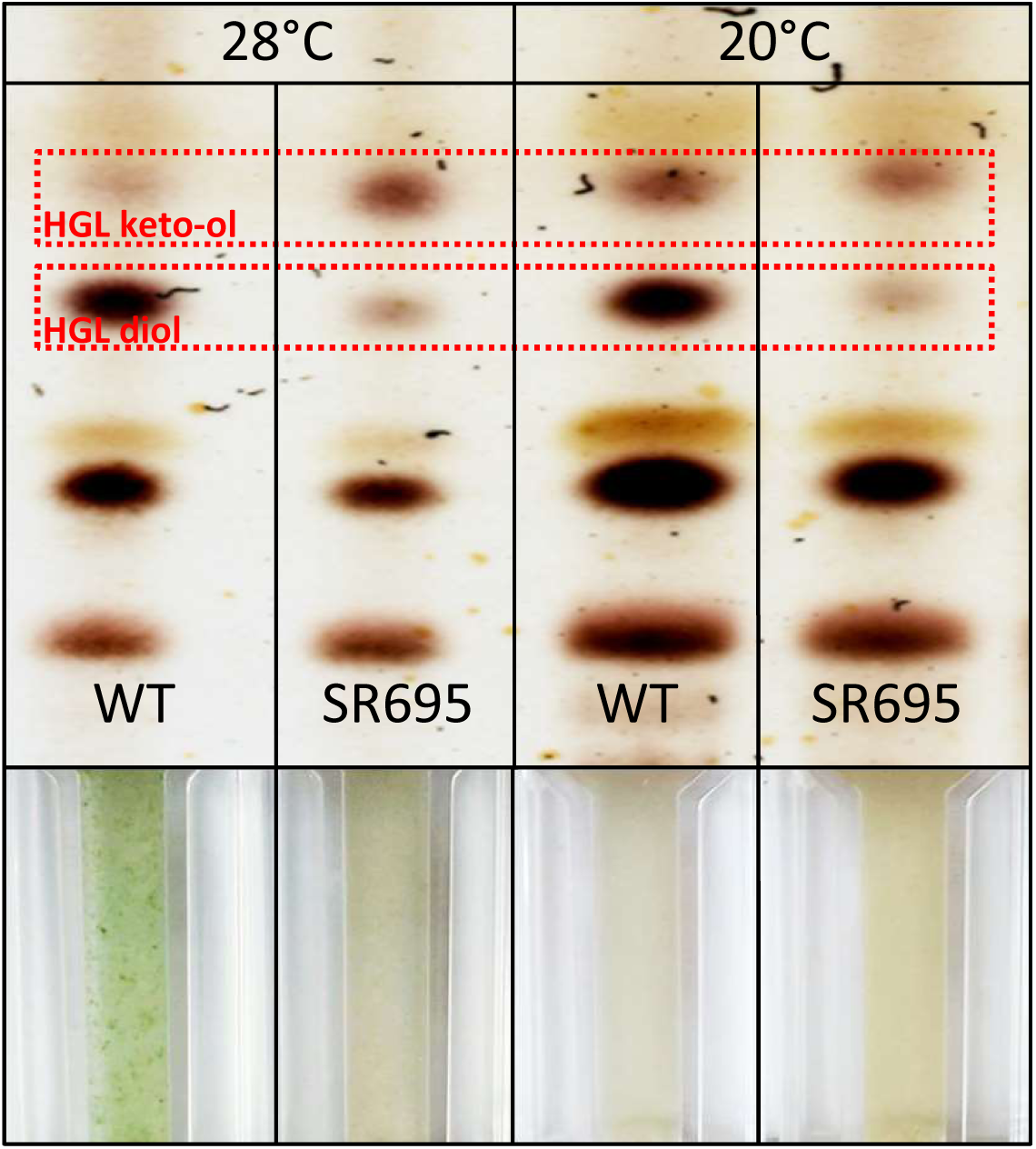
Comparison of the glycolipid composition of wild-type and mutant SR695 heterocysts. TLC analysis of lipids obtained from whole cell extracts of cultures incubated at the indicated temperatures. Red boxes, heterocyst-specific glycolipids; lower panels, photographs of the respective cultures.

### The protein HgdA is soluble and forms dimers *in vitro*

For the biochemical characterization of the HgdA protein, we overexpressed the gene in *Escherichia coli* and purified the protein by affinity chromatography, followed by size-exclusion chromatography. The major peak of HgdA in the size-exclusion chromatography elution profile corresponded to the dimeric form; additional peak shoulders, probably representing monomeric and other oligomeric forms of HgdA, were also present (Fig. 5A).

**Figure 5.**
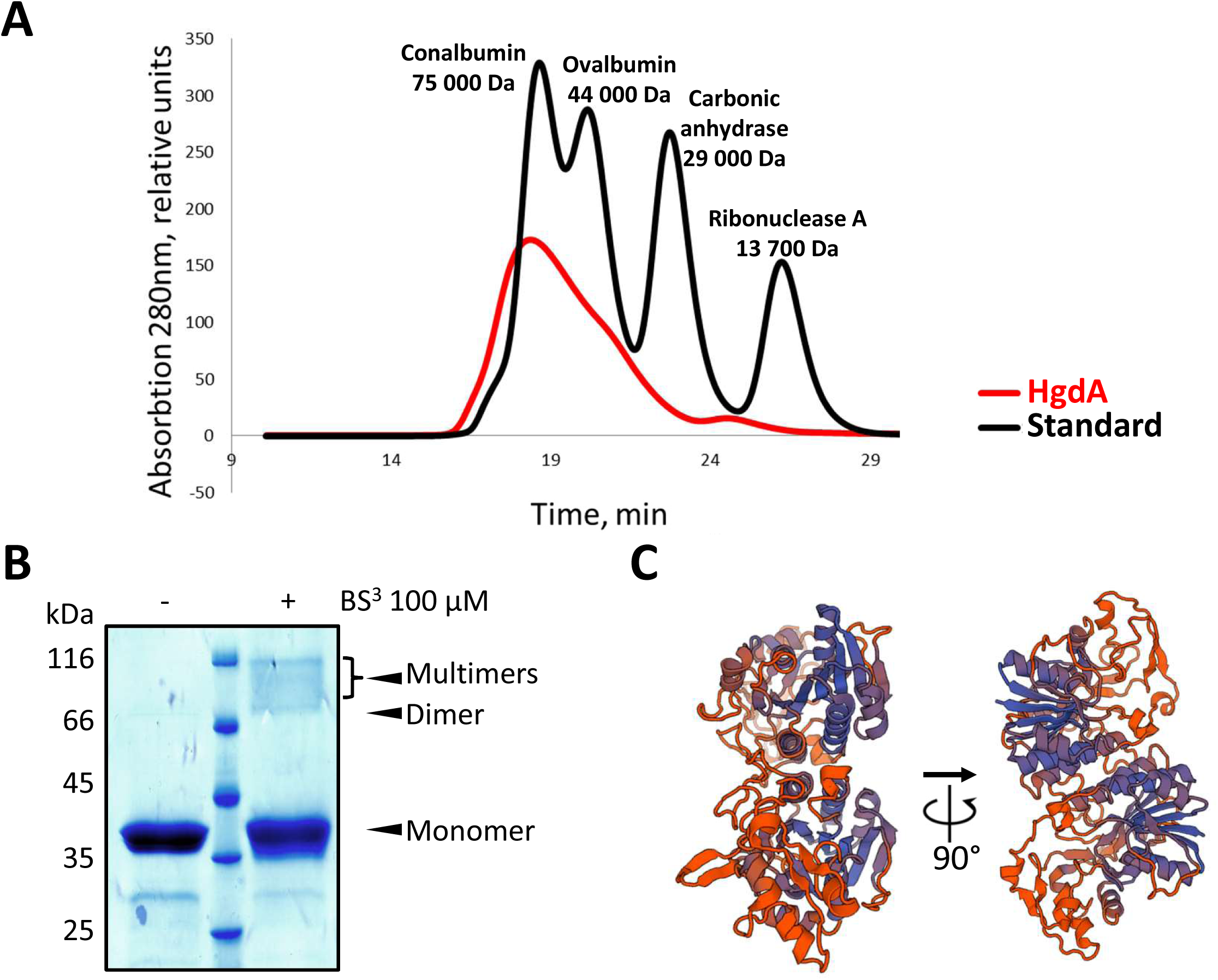
Analysis of the oligomeric state of HgdA. **A)** Size-exclusion chromatography elution profile of recombinant HgdA and a mixture of standard proteins. Peak fractions containing HgdA corresponds to the dimer. **B)** SDS-PAGE of recombinant HgdA cross-linked with BS^3^. **C)** Predicted structure of the HgdA dimer based on the structure of the homologous enzyme WbgU (a UDP-GalNAc 4-epimerase), created using a Swiss model online tool.

On SDS-polyacrylamide gels, the band of purified HgdA consisted of the monomeric form. However, when purified HgdA was incubated with the amino-reactive cross-linker suberic acid bis(3-sulfo-*N*-hydroxysuccinimide ester) (BS^3^), which forms covalent bonds between interacting proteins, also dimeric and other oligomeric forms were detected, and the band representing the monomeric form was less intense than the band in the sample without crosslinker (Fig. 5B).

We modeled the structure of HgdA using the Swiss model online tool (46) based on its closest homolog with a solved structure, namely the UDP-GalNAc 4-epimerase WbgU. The modeling revealed that HgdA probably forms dimers (Fig. 5C), in agreement with the results of size-exclusion chromatography and crosslinking experiments.

### HgdA fulfills a UDP-galactose 4-epimerase function *in vitro*

Based on the sequence similarity of HgdA to UDP-galactose 4-epimerase, we tested whether HgdA converts UDP-galactose to UDP-glucose (47). The enzyme catalyzed the conversion at a rate of approximately 30-40 nmol min^−1^ nmol HgdA^−1^ depending on the protein concentration. UDP-glucose production by HgdA increased when higher concentrations of the enzyme were used; the activity was considerably lower when tested at 99°C (Table 1, Fig. S3). UDP-galactose 4-epimerases use NAD as a cofactor, which is constantly bound in the conserved cofactor-binding glycine-rich site in the Rossmann fold (24, 27, 48, 49). However, we were unable to extract or detect NAD from the enzyme using standard protocols (50). In place of the conserved NAD-binding motif GXXGXXG, the HgdA sequence has a GIDEFIG motif, with the second glycine replaced by glutamate (Fig. S1). An NCBI BLAST search showed that such a motif is also found in HgdA homologs in several other cyanobacteria.

**Table 1.** Enzymatic activity of HgdA. Glucose oxidase-horseradish peroxidase assay of UDP-galactose 4-epimerase activity of recombinant HgdA. HgdA at the concentration of 10 µM was incubated with 1 mM UDP-galactose at indicated temperatures for 2 hours, and UDP-glucose production was measured. Bovine serum albumin (BSA) was used as negative control. Shown is the relative enzymatic activity in arbitrary units (AU) with the standard deviation of indicated experimental replicates.

|  | Relative activity, AU | Standard deviation, AU | # replicates |
| --- | --- | --- | --- |
| HgdA, reaction at 37°C | 1.196 | 0.061 | 6 |
| BSA, reaction at 37°C | 1.026 | 0.028 | 6 |
| HgdA, reaction at 99°C | 1.104 | 0.023 | 2 |
| BSA, reaction at 99°C | 1.035 | 0.014 | 2 |

## Discussion

One of the main events in heterocyst maturation is the formation of the heterocyst-specific envelope. A variety of enzymes participate in this process, including those that are responsible for the synthesis and transport of the envelope components (16, 18, 22, 23, 51–53). In this study, we investigated the function of the putative epimerase HgdA (Fig. 1) in heterocyst formation. Our results partially confirmed previous findings (22, 23), and in addition described the enzymatic activity of HgdA.

Transcripts of *hgdA* were found only when the heterocysts were almost completely mature (24–48 h after nitrogen stepdown; Fig. 2A). These time points were markedly later than activation of the *devB* gene, which encodes the membrane fusion component of the efflux pump transporting HGLs (16, 18). However, the expression patterns of the *hgdB* and *hgdC* genes (23) are similar to that of *hgdA*, which indicates that products of the *hgdBCA* gene cluster are formed at the same time even if they probably do not comprise an operon (22) and that the proteins might work together. The localization of HgdA in the cytoplasm of mature heterocysts (Fig. 2B, C) confirms its specific importance for these differentiated cells and demonstrates that HgdA is a soluble protein, as expected from the *in silico* analysis of its sequence.

Our mutant SR695, like the transposon-insertion mutant of this gene (22) showed a Fox^−^ phenotype, i.e., the inability to grow diazotrophically under oxic conditions. Since nitrogenase activity was detectable under anoxic conditions, this mutant shows a phenotype, which is specific for mutants with an impaired heterocyst envelope (Fig. 3A-C).

Although the hgl layer was present in the mutant SR695 (Fig. 3D), its defect allowed oxygen to enter the heterocyst. The main difference between the mutant and wild-type was in the HGL composition (Fig. 4). Specifically, the aberrant ratio of the two major HGLs in the mutant, with an excess of HGL_26_ diol, seemed to be critical for hgl layer formation and heterocyst function at 28 °C. The same aberrant HGL ratio, which causes a Fox^−^ phenotype, in an *hgdB* mutant has been found (23); this finding along with the results of our expression studies might indicate that products of the *hgdB, hgdC,* and *hgdA* genes work cooperatively. Most UDP-galactose 4-epimerases form dimers or other oligomers (24), but they can also function in a monomeric state (54). Our size-exclusion chromatography, crosslinking, and modeling results indicated that the main active states of HgdA are probably dimers (Fig. 5).

Purified HgdA has typical UDP-galactose 4-epimerase activity *in vitro* (Table 1). Compared to other known UDP-galactose 4-epimerase activities, this activity was in the lower range, with some epimerases having activities several times less and others hundreds of times more than that of HgdA (55–59). Since we were unable to detect or extract NAD from the purified active HgdA protein, we assume that the altered NAD-binding sequence, with glutamate replacing glycine, captures more tightly NAD.

Based on our results, we propose a function of HgdA in the HGL synthesis (Fig. 6). HgdA converts UDP-galactose derived from thylakoid degradation, which takes place during heterocyst differentiation, to UDP-glucose. According to our model, some UDP-galactose in the heterocyst is used for the production of the keto-ol form by a specific glycosyltransferase (#1), and some is utilized by HgdA to produce UDP-glucose. The produced UDP-glucose is then bound to the HGL_26_ diol aglycon by the other specific glycosyltransferase (#2). The *Anabaena* sp. genome carries a number of genes encoding glycosyltransferases, including several (*all5341, all5342, all5343*) close to the *hgdA* gene (22, 60).

**Figure 6.**
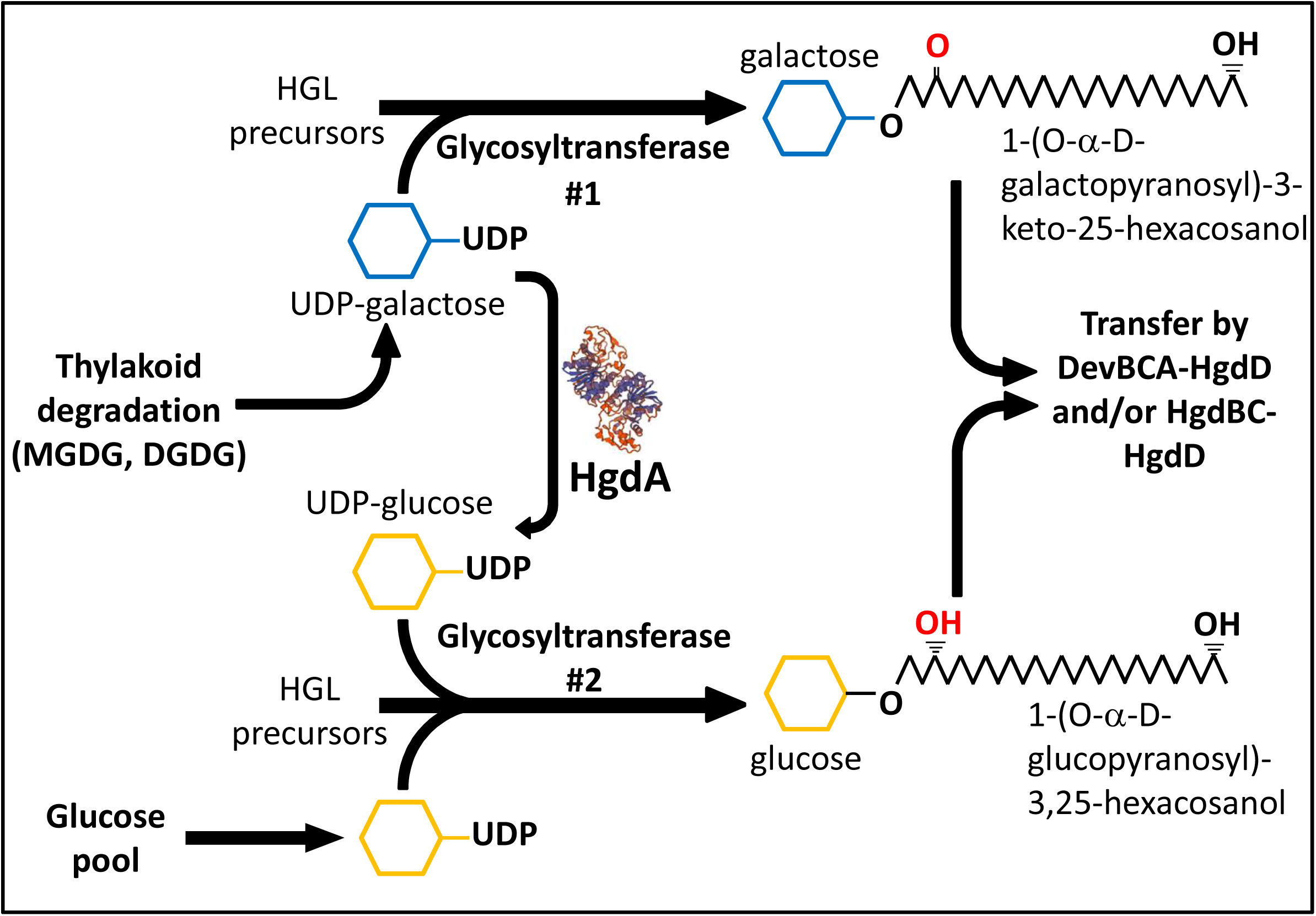
Proposed function of HgdA in HGL synthesis. HgdA is an epimerase that possibly epimerizes galactose, obtained from different sources including digalactosyldiacylglycerol (DGDG) and monogalactosyldiacylglycerol (MGDG), to glucose. Thereafter, different glycosyltransferases bind glucose or galactose to different aglycone precursors of HGLs.

In the *hgdA* mutant, insufficient amounts of the substrate UDP-galactose are present for sufficient HGL_26_ diol production, and in addition, residual UDP-galactose is utilized to synthesize other HGL keto-ols, which leads to the altered HGL diol:keto-ol ratio observed. However, other HGL diol biosynthetic pathways independent of HgdA must be present because mutant SR695 heterocysts contain a small amount of the HGL diol (Fig. 4), which nevertheless does not support heterocyst function. The similarity of the phenotypes of the *hgdA* mutant and *hgdB* mutant (23) suggest that HgdA and HgdBC closely cooperate, but further investigation is required.

Altered HGL diol:keto-ol ratios have been described in other situations. For instance, during growth at higher temperatures, cyanobacteria produce higher amounts of HGL diols (43, 44), which might protect heterocysts from gas penetration under these conditions. When HGL keto-ols are prevalent and the amount of HGL diols is lower, the heterocyst cell envelope might lose its gas tightness at higher temperatures; at lower temperatures, when amounts of the keto-ol form increase, the envelope retains its gas tightness.

The deposition of the HGLs in the wild-type and in the *hgdA* or *hgdB* mutant (22, 23) differ (Fig. 7). The wild-type forms a normal hgl layer around the entire heterocyst using two exporter systems, namely DevBCA-HgdD (18) mostly at the polar neck regions and HgdBC-HgdD (23) at the lateral sides. In the *hgdB* mutant, the hgl layer is replaced by an amorphous layer at the lateral sides because the HgdBC transporter is lacking, and the hgl layer is thicker at the polar regions because of excess substrate for DevBCA-HgdD (HGLs that are not transported by HgdBC-HgdD but are still synthesized). In the *hgdA* mutant, both transporters are present, but because HGL production is deficient, the hgl layer is much thinner than in the wild-type.

**Figure 7.**
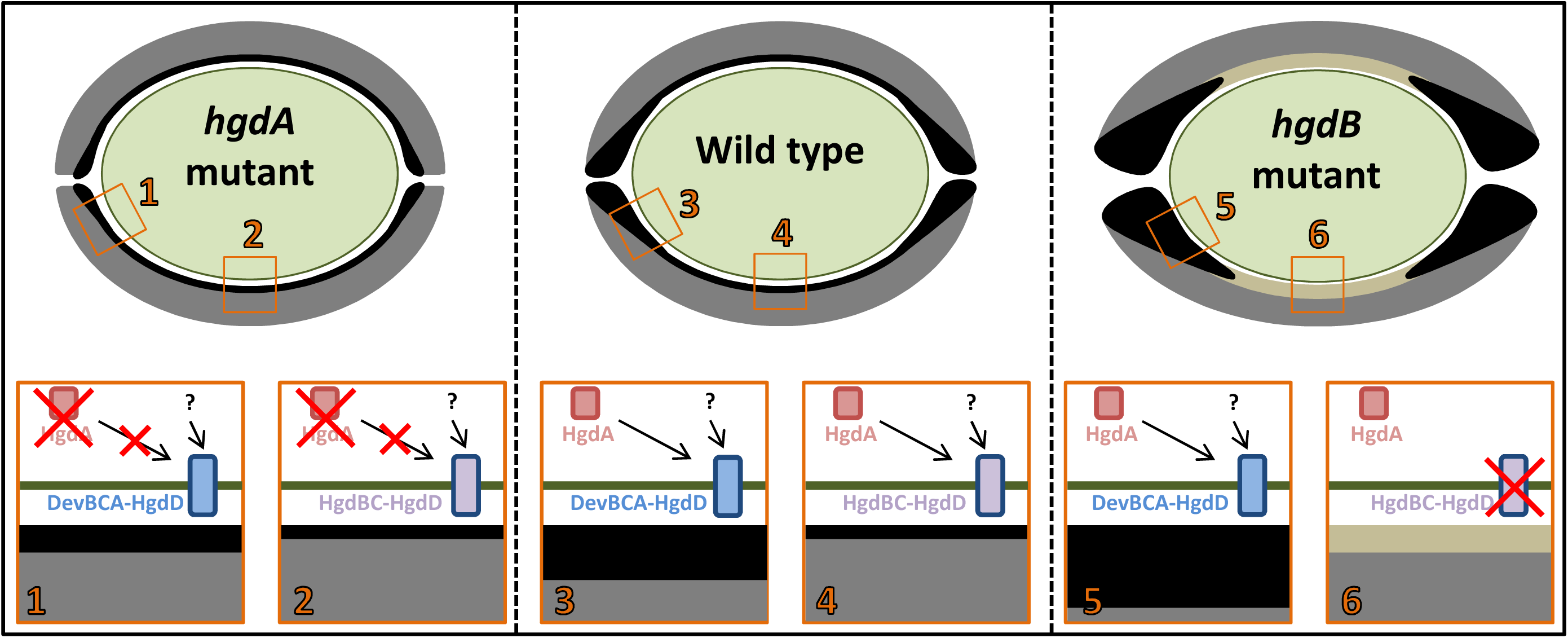
Comparison of hgl layers of heterocysts from wild-type, *hgdA* mutant, and *hgdB* mutant of *Anabaena* sp. The wild-type heterocyst possesses normal hgl (black) and hep (gray) layers. The *hgdA* mutant heterocyst has a much thinner hgl layer. The *hgdB* mutant heterocyst (22, 23) has a thicker hgl layer at the polar regions but it is replaced by an amorphous unstructured layer at the lateral sides of the cell. Green line, cellular membranes and cell wall. Lower panels show the putative interplay between the different means of HGL synthesis and transport involving HgdA and transporters DevBCA-HgdD (18) or HgdBC-HgdD (23). Arrows from HgdA show the route of HGLs, produced with help of HgdA, to a transporter (DevBCA-HgdD or HgdBC-HgdD). Arrows from the question marks indicate routes of HGLs produced independently of HgdA (probably at earlier stages of heterocyst development).

In conclusion, our results indicate that the epimerase HgdA takes part in the synthesis of the HGL diol form, thereby controlling the HGL keto-ol:diol ratio, and probably works at the late stages of heterocyst development and fine-tunes the proportion of HGL in the heterocyst envelope. At this stage, much UDP-galactose (HgdA substrate) must be available in the heterocysts; this substrate originates from the main components of thylakoid membranes, digalactosyldiacylglycerol (DGDG) and monogalactosyldiacylglycerol (MGDG) (61–63), which are degraded during heterocyst maturation. HgdA probably takes part in the synthesis of HGLs at some point, and a transporter composed of HgdBC and HgdD (TolC homologue) might export HgdA products directly or sequentially to the heterocyst cell envelope.

## Materials and Methods

### Organisms and growth conditions

*Anabaena* sp. PCC 7120 wild-type and its derivative mutant strains were cultivated in liquid BG11 medium (64) in 100 mL Erlenmeyer flasks under continuous illumination (17–22 μmol photons m^−2^ s^−1^) at 28 °C with shaking at 120 rpm. For RNA isolation, cells were cultivated in 700 mL of nitrate-free BG11 medium (BG11_0_) supplemented with 2.5 mM NH_4_Cl as nitrogen source and 5 mM TES buffer (pH 7.8) in one-liter bottles continuously supplied with CO_2_-enriched air (2%). Mutant strains were cultivated in BG11 medium supplemented with spectinomycin and streptomycin (2.5 µg mL^−1^ each).

For the nitrogen stepdown experiments, cells were washed three times in BG11_0_ medium and cultivated afterwards in BG11_0_.

All cloning and plasmid maintenance occurred in *E. coli* strains Top10, NEB10, Lemo21 (DE3), and HB101. For triparental mating, *E. coli* strain J53 (bearing the conjugative plasmid RP4), strain HB101 (bearing the helper plasmid pRL528 and the cargo plasmid pRL277 with a fragment of the gene of interest), and wild-type *Anabaena* sp. were used (65–67) (Tables S1).

The *hgdA* gene for protein synthesis was overexpressed in *E. coli* Lemo21 (DE3) (Table S1).

### DNA manipulations

To construct an insertion mutant of *hgdA* by homologous recombination, an internal fragment of the gene was amplified by PCR (see Table S2 for primers) with 1 µL of the wild-type *Anabaena* sp. culture as template and cloned into the *Xho*I-restricted suicide vector pRL277 (Table S1) using Gibson assembly (68) (Fig. S2A). The resulting plasmid pIM695 was transferred into wild-type *Anabaena* sp. cells by triparental mating, followed by selection on streptomycin- and spectinomycin-containing BG11 agar plates. In the antibiotic-resistant *Anabaena* sp. colonies, where a single recombination event between the *hgdA* gene in the genome and its internal fragment in the pIM695 vector had occurred, the *hgdA* gene was disrupted by the pRL277 vector (Fig. 1A, S2A). Full segregation of one selected mutant (SR695) colony was confirmed by PCR (Fig. S2B) with a small piece of the mutant colony as template.

To localize HgdA in *Anabaena* sp. filaments, a plasmid with a translational fusion of the HgdA C-terminus with the super-folder GFP (sfGFP) (69) was constructed following the method described in (23). The 3′-end of *hgdA* and the entire *sfGFP* were amplified by PCR and cloned into the *Xho*I-restricted suicide vector pRL277 using Gibson assembly. The resulting plasmid pIM717 was transferred into wild-type *Anabaena* sp. cells using triparental mating, followed by positive colony selection on streptomycin- and spectinomycin-containing BG11 agar plates. *Anabaena* sp. colonies contained the *hgdA* gene fused with *sfGFP* (strain SR717). The fusion was confirmed by PCR.

For complementation of the SR695 mutant, the *hgdA* gene under control of the *glnA* promoter (37) was cloned into the *EcoR*I-restricted self-replicating plasmid pIM612, which bears a neomycin-resistance cassette (36), using Gibson assembly. The resulting plasmid pIM774 was transferred into mutant SR695 cells, and positive colonies were selected on BG11 agar plates containing neomycin, streptomycin, and spectinomycin. The presence of the undisrupted *hgdA* gene in the complemented mutant colonies was confirmed by PCR (Fig. S2B).

For overexpression of the *hgdA* gene in *E. coli, hgdA*, followed by sequences encoding a Strep-tag and His-tag at the 3′-terminus was cloned into plasmid pET15b (Novagen, Merck) digested with *Nco*I, with help of Gibson assembly to yield plasmid pIM753.

### RNA isolation and RT-PCR

RNA was isolated at different time points after nitrogen stepdown using UPzol reagent (Biotechrabbit, Henningsdorf) according to the manufacturer’s instructions from wild-type *Anabaena* sp. cells grown in bottles as described above. The purity and concentration of the extracted RNA were estimated by electrophoresis and GelQuantNET software (biochemlabsolutions.com). Reverse transcription (RT) reactions were performed using the Applied Biosystems RT-reaction kit. The primers used for all PCR reactions are listed in Table S2.

### Microscopy

For light and fluorescence microscopy, wild-type and mutant *Anabaena* sp. cells were placed onto agarose-covered glass slides and observed under a Leica DM 2500 microscope connected to a Leica DFC420C camera or a Leica DM5500 B microscope connected to a Leica monochrome DFC360 FX camera.

For electron microscopy, cells were fixed and post-fixed with glutaraldehyde and potassium permanganate respectively (16). Ultrathin sections were stained with uranyl acetate and lead citrate and examined with a Philips Tecnai 10 electron microscope at 80 kHz.

### Staining methods for light microscopy

Cells were stained with BODIPY (boron dipyrromethene difluoride 493/503, Molecular Probes) following the protocol described by Perez et al. (41). Briefly, 1 mL of *Anabaena* sp. cell suspension was centrifuged at 4000 × *g* for 10 min, washed with PBS buffer, and resuspended in 200 µL PBS. BODIPY (1 μL of 50 ng mL^−1^ in DMSO) was added. The cell suspension was incubated in the dark for 30 min at room temperature and examined by light and fluorescence microscopy. Fluorescence or phase-contrast images were captured with a Leica DM 5500B microscope connected to a Leica monochrome DFC360 FX camera.

Cells were stained with alcian blue following the protocol described in (42). Cell suspensions were mixed with 1.5% alcian blue in water (at a ratio of 20:1) and incubated at room temperature for 5 min. For triphenyl tetrazolium chloride (TTC) staining, cell suspensions were mixed with TTC solution (0.01% TTC, w/v, in the final mixture) and incubated in the dark for 10 min at room temperature (38). Filaments stained with TTC or alcian blue were examined using a Leica DM 2500 microscope connected to a Leica DFC420C camera.

### Analysis of heterocyst-specific glycolipids

Glycolipids were analyzed by thin-layer chromatography as described in (70) with minor modifications. In brief, wild-type and mutant cells of equal chlorophyll *a* concentration [measured according to (71)] were pelleted and resuspended in methanol-chloroform (1:1). The solvents were evaporated under air in a fume hood. Lipids were dissolved in chloroform and applied to a silica-gel-coated aluminum plate (Macherey-Nagel, #818033). Thin-layer chromatographs were run with a mobile phase composed of chloroform:methanol:acetic acid:water (23:4:2.7:1). Lipids were visualized by spraying the plate with 25–50% sulfuric acid and exposing it to 180 °C for 60–120 s.

### Nitrogenase activity

Nitrogenase activity was measured using the acetylene reduction method for cyanobacteria (40). Briefly, cultures were incubated in the presence of acetylene for several hours in flasks closed with gas-tight caps. Anoxic conditions were generated before incubation with acetylene by adding 3-(3,4-dichlorophenyl)-1,1-dimethylurea (DCMU, 10 µM, in methanol); the sealed flasks were then filled with argon and incubated for 1 h. For oxic conditions, this step was omitted. After incubation with acetylene, 1 mL of the gaseous phase was taken from each flask, and the amount of ethylene produced was measured by gas chromatography.

### Preparation of *Anabaena* sp. vegetative and heterocyst cell lysates

Heterocysts were isolated as previously described (72, 73). Briefly, after nitrogen stepdown and incubation for 3 days, cells were collected by centrifugation; the pellet was resuspended in 15 ml of ice-cold 8% sucrose, 5% Triton X-100, 50 mM EDTA pH 8.0, 50 mM Tris pH 8.0, and 1 mg/ml of lysozyme. The suspensions were mixed vigorously on a vortex shaker for 2–3 min at room temperature. The solution was mildly sonicated with a Branson sonifier (3 × 3 min, 30% duty cycle, 3 output control). Heterocysts were collected by centrifugation at 3000 × *g* for 5 min at 4 °C; the supernatant was the vegetative cell lysate. Heterocysts were washed several times in 8% sucrose, 50 mM EDTA pH 8.0, 50 mM Tris pH 8.0.

To obtain the soluble (cytoplasmic) heterocyst fraction, the heterocyst pellet was resuspended in 5 mM HEPES buffer (pH 8.0) containing 1 mM phenylmethylsulfonyl fluoride (PMSF). The suspension was strongly sonicated (5 × 3 min, 50% duty cycle, 5 output control). The cells were then passed through a French pressure cell (SLM instruments, Inc) at 1100 Psi 4– 5 times. The suspension was centrifuged at 3000 × *g* for 30 min at 4 °C to separate undisrupted heterocysts. Then the supernatant was centrifuged at 15,000 × *g* for 1 h at 4 °C. The supernatant of this last centrifugation step contained the heterocyst cytoplasmic fraction; the pellet consisted of insoluble debris and membranes.

Samples were analyzed by Western blotting with polyclonal antibodies raised against the peptide synthesized from the C-terminus of HgdA (NH_2_-CQTKNWLQNTDIQKLVK-COOH). Peptides were synthesized and antibodies were produced by Pineda Antibody-service (Berlin). Rabbit polyclonal antibodies raised against the PII protein of *Synechococcus* sp. (74) were used as an internal control. After incubation with anti-HgdA and anti-PII antibodies, the membranes were washed in PBS buffer containing 0.05% Tween 20 (Carl Roth) and incubated with secondary peroxidase-coupled anti-rabbit IgG antibodies (Sigma A6154). For detection, Lumi-Light western blotting substrate (Roche) and a Gel Logic 1500 imager (Kodak) were used.

### Overexpression of *hgdA* and purification of HgdA

The *hgdA* gene was overexpressed in *E. coli* Lemo21 (DE3) cells carrying the pIM753 plasmid. Cells were cultivated in 3 L Erlenmeyer flasks at 37 °C with continuous shaking at 120 rpm until they reached an OD_600_ of 0.6. Gene expression was induced by adding isopropyl β-D-1-thiogalactopyranoside (Carl Roth) at a final concentration of 0.1 mM and incubation of the flasks at 25 °C overnight with shaking. After induction, cells were pelleted at 7000 × *g* for 15 min at 4 °C, and the pellet was resuspended in lysis buffer (20 mM Tris, 200 mM NaCl, 0.5% Triton X-100, pH 7.5) containing 1 mM PMSF and 1 mg/mL lysozyme and incubated at room temperature for 1–2 h. Then, the solutions were sonicated with a Branson sonifier (3 × 3 min, 50% duty cycle, 5 output control) and centrifuged at 17,000 × *g* for 30 min at 4 °C. The supernatant, which contained extracted soluble proteins, was used for purification of HgdA by affinity chromatography using a Strep-column (IBA-Lifesciences) and Tris buffer (20 mM Tris, 200 mM NaCl, pH 7.5) for equilibration of the column and washing steps; the same buffer containing 2.5 mM desthiobiotin was used for elution. The eluted fractions were pooled and concentrated, and the purity of the HgdA protein was checked by SDS-PAGE.

HgdA was more highly purified and its oligomeric state was estimated by size-exclusion chromatography using an ÄKTA chromatography system and a Superdex 75 10/300 column in Tris buffer (see above). To calculate the molecular masses of the proteins in the eluted peaks, a mixture of standard proteins (Gel Filtration LMW Calibration Kit, GE Life Sciences) was run through the column. The fractions corresponding to different peaks of HgdA purification were pooled, concentrated, and analyzed by SDS-PAGE. The concentration of pure HgdA protein was determined by the Bradford method using Roti-Quant solution (Carl Roth).

### Crosslinking assay

Interacting proteins were crosslinked with suberic acid bis(3-sulfo-*N*-hydroxysuccinimide ester) (BS^3^), which crosslinks epimerases (75). Purified HgdA (5 µM) was incubated with BS^3^ (100 µM) for 30 min at 37 °C in 25 µL of Tris buffer (see above). Afterwards, the entire sample was used for SDS-PAGE analysis.

### Epimerase activity assay

To test the epimerase activity of HgdA, an established colorimetric glucose oxidase-horseradish peroxidase (GOD-POD) coupled assay was used (47, 59, 76). In brief, 1 mM UDP-Gal dissolved in 20 mM Tris buffer containing 200 mM NaCl, pH 7.5 was incubated with different amounts of purified HgdA in a total reaction volume of 22 μL at 37 °C for 1 or 2h. Reactions were stopped and proteins were acid-hydrolyzed with 3.5 μL of 0.4 N HCl at 100 °C for 6 min. The mixture was neutralized with 3.5 μl of 0.4 N NaOH. Aliquots (7.5 μL) were taken from each reaction mixture and applied to a 96-well plate. The GOD-POD assay was used to detect released glucose according to the manufacturer’s instructions (Sigma-Aldrich). The reaction was stopped and the color was developed by adding 100 μL of 6 N HCl per well. Afterwards, the absorbance at 540 nm was read by a TECAN Spark 10M plate reader.

## Acknowledgments

We thank Claudia Menzel for electron microscopy sample preparation, Thomas Härtner for help with gas chromatography, Oliver Betz for access to the TEM, Ritu Garg for help with size exclusion chromatography of standard proteins, and Karl Forchhammer for fruitful discussions and valuable advice. We also thank Karen Brune for critically reading the manuscript and improving the text linguistically. This work was supported by Deutsche Forschungsgemeinschaft (SFB766 and GRK1703).

## Figure legends

**Figure S1. COBALT multiple alignment of several characterized HgdA homologs found using the online tool PaperBLAST.** Red, highly conserved residues; blue, less conserved residues; gray, not conserved residues; yellow box, conserved GXXGXXG NAD(P) binding site; violet boxes, residues of the conserved S(X)_24_–Y(X)_3_K catalytic triad.

**Figure S2. Construction and segregation of the *hgdA* mutant.**

**A)** Construction of an *hgdA* mutant via homologous recombination. Arrows with numbers, primers used for the genotypic analysis of the mutants (see also Table S2). **B)** Segregation of the *hgdA* mutant analyzed by PCR. SR695, mutant SR695; wt, wild-type; SR695c, SR695 mutant complemented with the *hgdA* gene. Primer numbers correspond to those depicted in **A**.

**Figure S3. Enzymatic activity of HgdA.**

**A)** and **B)** Glucose oxidase-horseradish peroxidase assay of UDP-galactose 4-epimerase activity of recombinant dimeric HgdA. HgdA (red) at the indicated concentrations was incubated at 37 °C **(A)** or at concentration 10 µM at 37 °C or 99 °C **(B)** with 1 mM UDP-galactose, and UDP-glucose production was measured (expressed in units of absorbance at 540 nm, Abs_540_). Each point represents the mean value ± standard deviation of two experimental replicates. Bovine serum albumin (BSA) was used as negative control (black). In **A**, a representative of two independent experiments is shown.

## References

1. Adams DG, Duggan PS. 1999. Tansley Review No. 107 Heterocyst and akinete differentiation in cyanobacteria. New Phytol 144:3–33.

2. Maldener I, Summers ML, Sukenik A. 2014. Cellular differentiation in filamentous cyanobacteria., p. 263–291. *In* Flores, E, Herrero, A (eds.), The Cell Biology of Cyanobacteria. Caister Academic Press, Norfolk, UK.

3. Kumar K, Mella-Herrera RA, Golden JW. 2010. Cyanobacterial heterocysts. Cold Spring Harb Perspect Biol 2:a000315.

4. Muro-Pastor AM, Hess WR. 2012. Heterocyst differentiation: From single mutants to global approaches. Trends Microbiol 20:548–557.

5. Walsby AE. 2007. Cyanobacterial heterocysts: terminal pores proposed as sites of gas exchange. Trends Microbiol 15:340–349.

6. Gambacorta A, Pagnotta E, Romano I, Sodano G, Trincone A. 1998. Heterocyst Glycilipids From Nitrogen-Fixing Cyanobacteria Other Than Nostocaceae. Phytochemistry 48:801–805.

7. Soriente A, Gambacorta A, Trincone A, Sili C, Vincenzini M, Sodano G. 1993. Heterocyst glycolipids of the cyanobacterium *Cyanospira rippkae*. Phytochemistry 33:393–396.

8. Gambacorta A, Trincone A, Soriente A, Sodano G. 1999. Chemistry of glycolipids from the heterocysts of nitrogen-fixing cyanobacteria. Curr Top Phytochem 2:145–150.

9. Bauersachs T, Compaoré J, Hopmans EC, Stal LJ, Schouten S, Sinninghe JS. 2009. Distribution of heterocyst glycolipids in cyanobacteria. Phytochemistry 70:2034–2039.

10. Schouten S, Villareal TA, Hopmans EC, Mets A, Swanson KM, Damste JSS. 2013. Endosymbiotic heterocystous cyanobacteria synthesize different heterocyst glycolipids than free-living heterocystous cyanobacteria. Phytochemistry 85:115–121.

11. Bale NJ, Hopmans EC, Zell C, Sobrinho RL, Kim J-H, Damste JSS, Villareal TA, Schouten S. 2015. Long chain glycolipids with pentose head groups as biomarkers for marine endosymbiotic heterocystous cyanobacteria. Org Geochem 81:1–7.

12. Bale NJ, Hopmans EC, Dorhout D, Stal LJ, Grego M, van Bleijswijk J, Sinninghe Damsté JS, Schouten S. 2018. A novel heterocyst glycolipid detected in a pelagic N_2_-fixing cyanobacterium of the genus Calothrix. Org Geochem.

13. Gambacorta A, Romano I, Trincone A, Soriente A, Giordano M, Sodano G. 1996. Heterocyst glycolipids from five nitrogen-fixing cyanobacteria. Gazz Chim Ital 126:653–656.

14. Awai K, Lechno-Yossef S, Wolk CP. 2009. Heterocyst Envelope Glycolipids, p. 179–202. *In* Wada, H, Murata, N (eds.), Lipids in Photosynthesis: Essential and Regulatory Functions. Springer, Dordrecht.

15. Maldener I, Fiedler G, Ernst A, Fernândez-Pinas F, Wolk CP. 1994. Characterization of *devA,* a Gene Required for the Maturation of Proheterocysts in the Cyanobacterium *Anabaena* sp. Strain PCC 7120. J Bacteriol 176:7543–7549.

16. Fiedler G, Arnold M, Hannus S, Maldener I. 1998. The DevBCA exporter is essential for envelope formation in heterocysts of the cyanobacterium *Anabaena* sp. strain PCC 7120. Mol Microbiol 27:1193–1202.

17. Staron P, Forchhammer K, Maldener I. 2014. Structure-function analysis of the ATP-driven glycolipid efflux pump DevBCA reveals complex organization with TolC/HgdD. FEBS Lett 588:395–400.

18. Staron P, Forchhammer K, Maldener I. 2011. Novel ATP-driven pathway of glycolipid export involving TolC protein. J Biol Chem 286:38202–10.

19. Staron P. 2012. PhD thesis. Structural and Functional Characterization of the ATP-Driven Glycolipid-Efflux Pump DevBCA-TolC and its Homologues in the Filamentous Cyanobacterium Anabaena sp. PCC 7120. Eberhard-Karls Universität Tübingen, Tübingen.

20. Shvarev D, Maldener I. 2018. ATP-binding cassette transporters of the multicellular cyanobacterium *Anabaena* sp. PCC 7120: a wide variety for a complex lifestyle. FEMS Microbiol Lett 365:1–6.

21. Staron P, Maldener I. 2012. All0809/8/7 is a DevBCA-like ABC-type efflux pump required for diazotrophic growth in *Anabaena* sp. PCC 7120. Microbiology 158:2537–45.

22. Fan Q, Huang G, Lechno-Yossef S, Wolk CP, Kaneko T, Tabata S. 2005. Clustered genes required for synthesis and deposition of envelope glycolipids in *Anabaena* sp. strain PCC 7120. Mol Microbiol 58:227–43.

23. Shvarev D, Nishi C, Wörmer L, Maldener I. 2018. The ABC Transporter Components HgdB and HgdC are Important for Glycolipid Layer Composition and Function of Heterocysts in *Anabaena* sp. PCC 7120. Life 8:26.

24. Allard ST, Giraud MF, Naismith JH. 2001. Epimerases: structure, function and mechanism. Cell Mol Life Sci 58:1650–1665.

25. Maxwell ES. 1957. The enzymic interconversion of uridine diphosphogalactose and uridine diphosphoglucose. J Biol Chem 229:139–151.

26. Wilson DB, Hogness DS. 1964. The Enzymes of the Galactose Operon in Escherichia coli: I. Purification and Characterization of Uridine Diphosphogalactose 4-Epimerase. J Biol Chem 239:2469–2482.

27. Beerens K, Soetaert W, Desmet T. 2015. UDP-hexose 4-epimerases: a view on structure, mechanism and substrate specificity. Carbohydr Res 414:8–14.

28. Bauer AJ, Rayment I, Frey PA, Holden HM. 1992. The molecular structure of UDP-galactose 4-epimerase from *Escherichia coli* determined at 2.5 Å resolution. Proteins Struct Funct Bioinforma 12:372–381.

29. Giraud MF, Leonard GA, Field RA, Berlind C, Naismith JH. 2000. RmIc, the third enzyme of dTDP-L-rhamnose pathway, is a new class of epimerase. Nat Struct Biol 7:398–402.

30. Deacon AM, Ni YS, Coleman WG, Ealick SE. 2000. The crystal structure of ADP-L-glycero-D-mannoheptose 6-epimerase: Catalysis with a twist. Structure 8:453–462.

31. Carbone V, Schofield LR, Sang C, Sutherland-Smith AJ, Ronimus RS. 2018. Structural determination of archaeal UDP-N-acetylglucosamine 4-epimerase from *Methanobrevibacter ruminantium* M1 in complex with the bacterial cell wall intermediate UDP- N-acetylmuramic acid. Proteins Struct Funct Bioinforma 1–26.

32. Price MN, Arkin AP. 2017. PaperBLAST: Text Mining Papers for Information about Homologs. mSystems 2:e00039–17.

33. Dereeper A, Guignon V, Blanc G, Audic S, Buffet S, Chevenet F, Dufayard JF, Guindon S, Lefort V, Lescot M, Claverie JM, Gascuel O. 2008. Phylogeny.fr: robust phylogenetic analysis for the non-specialist. Nucleic Acids Res 36:465–469.

34. Dereeper A, Audic S, Claverie JM, Blanc G. 2010. BLAST-EXPLORER helps you build datasets for phylogenetic analysis. Evol Biol 10:8–13.

35. Paz-Yepes J, Flores E, Herrero A. 2009. Expression and Mutational Analysis of the glnB Genomic Region in the Heterocyst-Forming Cyanobacterium *Anabaena* sp. Strain PCC 7120. J Bacteriol 191:2353–2361.

36. Bornikoel AJ. 2018. PhD thesis. Key players in cell wall modification, multicellularity and cell-cell communication in the filamentous cyanobacterium Anabaena sp. PCC 7120. Eberhard-Karls Universität Tübingen, Tübingen.

37. Valladares A, Muro-pastor AM, Herrero A, Flores E. 2004. The NtcA-Dependent P1 Promoter Is Utilized for glnA Expression in N2-Fixing Heterocysts of *Anabaena* sp. Strain PCC 7120. J Bacteriol 186:7337–7343.

38. Fay P, Kulasooriya SA. 1972. Tetrazolium reduction and nitrogenase activity in heterocystous blue-green algae. Arch Mikrobiol 87:341–352.

39. Ernst A, Black T, Cai Y, Panoff J, Tiwari DN, Wolk CP. 1992. Synthesis of Nitrogenase in Mutants of the Cyanobacterium *Anabaena* sp. Strain PCC 7120 Affected in Heterocyst Development or Metabolism. J Bacteriol 174:6025–6032.

40. Bornikoel J, Staiger J, Madlung J, Forchhammer K, Maldener I. 2018. LytM factor Alr3353 affects filament morphology and cell-cell communication in the multicellular cyanobacterium *Anabaena* sp. PCC 7120. Mol Microbiol 108:187–203.

41. Perez R, Forchhammer K, Salerno G, Maldener I. 2016. Clear differences in metabolic and morphological adaptations of akinetes of two nostocales living in different habitats. Microbiol 162:214–223.

42. McKinney RE. 1953. Staining bacterial polysaccharides. J Bacteriol 66:453–454.

43. Bauersachs T, Stal LJ, Grego M, Schwark L. 2014. Temperature induced changes in the heterocyst glycolipid composition of N2 fixing heterocystous cyanobacteria. Org Geochem 69:98–105.

44. Wörmer L, Cires S, Velazquez D, Quesada A, Hinrichs K-U. 2012. Cyanobacterial heterocyst glycolipids in cultures and environmental samples: Diversity and biomarker potential. Limnol Oceanogr 57:1775–1788.

45. Perez R, Wörmer L, Sass P, Maldener I. 2017. A highly asynchronous developmental program triggered during germination of dormant akinetes of the filamentous diazotrophic cyanobacteria. FEMS Microbiol Ecol 94:10.1093/femsec/fix131.

46. Waterhouse A, Bertoni M, Bienert S, Studer G, Tauriello G, Gumienny R, Heer FT, De Beer TAP, Rempfer C, Bordoli L, Lepore R, Schwede T. 2018. SWISS-MODEL: Homology modelling of protein structures and complexes. Nucleic Acids Res 46:W296– W303.

47. Moreno F, Rodicio R, Herrero P. 1981. A new colorimetric assay for UDP-glucose 4-epimerase activity. Cell Mol Biol Incl Cyto Enzymol 27:589–592.

48. Rossmann MG, Moras D, Olsen KW. 1974. Chemical and biological evolution of a nucleotide-binding protein. Nature 250:194–199.

49. Bellamacina CR. 1996. The nicotinanamide dinucleotide binding motif: comparison of nucleotide binding proteins. FASEB J 10:1257–1269.

50. Creuzenet C, Belanger M, Wakarchuk WW, Lam JS. 2000. Expression, purification, and biochemical characterization of WbpP, a new UDP-GlcNAc C4 epimerase from *Pseudomonas aeruginosa* serotype O6. J Biol Chem 275:19060–19067.

51. Maldener I, Hannus S, Kammerer M. 2003. Description of five mutants of the cyanobacterium *Anabaena* sp. strain PCC 7120 affected in heterocyst differentiation and identification of the transposon-tagged genes. FEMS Microbiol Lett 224:205–213.

52. Huang G, Fan Q, Lechno-Yossef S, Wojciuch E, Wolk CP, Kaneko T, Tabata S. 2005. Clustered Genes Required for the Synthesis of Heterocyst Envelope Polysaccharide in *Anabaena* sp. Strain PCC 7120. J Bacteriol 187:1114–1123.

53. Nicolaisen K, Hahn A, Schleiff E. 2009. The cell wall in heterocyst formation by *Anabaena* sp. PCC 7120. J Basic Microbiol 49:5–24.

54. Nayar S, Bhattacharyya D. 1997. UDP-galactose 4-epimerase from *Escherichia coli:* Existence of a catalytic monomer. FEBS Lett 409:449–451.

55. Agarwal S, Gopal K, Upadhyaya T, Dixit A. 2007. Biochemical and functional characterization of UDP-galactose 4-epimerase from *Aeromonas hydrophila*. Biochim Biophys Acta - Proteins Proteomics 1774:828–837.

56. Chung SK, Ryu SI, Lee SB. 2012. Characterization of UDP-glucose 4-epimerase from *Pyrococcus horikoshii:* Regeneration of UDP to produce UDP-galactose using two-enzyme system with trehalose. Bioresour Technol 110:423–429.

57. Shin SM, Choi JM, Di Luccio E, Lee YJ, Lee SJ, Lee SJ, Lee SH, Lee DW. 2015. The structural basis of substrate promiscuity in UDP-hexose 4-epimerase from the hyperthermophilic Eubacterium *Thermotoga maritima*. Arch Biochem Biophys 585:39–51.

58. Guevara DR, El-Kereamy A, Yaish MW, Mei-Bi Y, Rothstein SJ. 2014. Functional characterization of the rice UDP-glucose 4-epimerase 1, OsUGE1: A potential role in cell wall carbohydrate partitioning during limiting nitrogen conditions. PLoS One 9:e96158.

59. Pardeshi P, Rao KK, Balaji P V. 2017. Rv3634c from *Mycobacterium tuberculosis* H37Rv encodes an enzyme with UDP-Gal/Glc and UDP-GalNAc 4-epimerase activities. PLoS One 12:e0175193.

60. Awai K, Wolk CP. 2007. Identifcation of the glycosyl transferase required for synthesis of the principal glycolipid characteristic of heterocysts of *Anabaena* sp. strain PCC 7120. FEMS Microbiol Lett 266:98–102.

61. Boudière L, Michaud M, Petroutsos D, Rébeillé F, Falconet D, Bastien O, Roy S, Finazzi G, Rolland N, Jouhet J, Block MA, Maréchal E. 2014. Glycerolipids in photosynthesis: Composition, synthesis and trafficking. Biochim Biophys Acta - Bioenerg 1837:470–480.

62. Maida E, Awai K. 2016. Digalactosyldiacylglycerol is essential in *Synechococcus elongatus* PCC 7942, but its function does not depend on its biosynthetic pathway. Biochim Biophys Acta - Mol Cell Biol Lipids 1861:1309–1314.

63. Yuzawa Y, Shimojima M, Sato R, Mizusawa N, Ikeda K, Suzuki M, Iwai M, Hori K, Wada H, Masuda S, Ohta H. 2014. Cyanobacterial monogalactosyldiacylglycerol-synthesis pathway is involved in normal unsaturation of galactolipids and low-temperature adaptation of *Synechocystis* sp. PCC 6803. Biochim Biophys Acta - Mol Cell Biol Lipids 1841:475–483.

64. Rippka R, Deruelles J, Waterbury JB, Herdman M, Stanier RY. 1979. Generic Assignments, Strain Histories and Properties of Pure Cultures of Cyanobacteria. Microbiology 111:1–61.

65. Wolk CP, Vonshak A, Kehoe P, Elhai J. 1984. Construction of shuttle vectors capable of conjugative transfer from Escherichia coli to nitrogen-fixing filamentous cyanobacteria. Proc Natl Acad Sci U S A 81:1561–1565.

66. Elhai J, Wolk CP. 1988. Conjugal Transfer of DNA to Cyanobacteria. Methods Enzymol 167:747–754.

67. Black TA, Cai Y, Wolk CP. 1993. Spatial expression and autoregulation of *hetR,* a gene involved in the control of heterocyst development in *Anabaena*. Mol Microbiol 9:77–84.

68. Gibson DG, Young L, Chuang R-Y, Venter JC, Hutchison CA, Smith HO. 2009. Enzymatic assembly of DNA molecules up to several hundred kilobases. Nat Methods 6:343–345.

69. Pédelacq J-D, Cabantous S, Tran T, Terwilliger TC, Waldo GS. 2006. Engineering and characterization of a superfolder green fluorescent protein. Nat Biotechnol 24:79–88.

70. Winkenbach F, Wolk CP, Jost M. 1972. Lipids of membranes and of the cell envelope in heterocysts of a blue-green alga. Planta 107:69–80.

71. Mackinney G. 1941. Absorption of Light by Chlorophyll Solutions. J Biol Chem 140:315–322.

72. Moslavac S, Reisinger V, Berg M, Mirus O, Vosyka O, Plöscher M, Flores E, Eichacker LA, Schleiff E. 2007. The proteome of the heterocyst cell wall in *Anabaena* sp. PCC 7120. Biol Chem 388:823–829.

73. Golden JW, Robinson SJ, Haselkorn R. 1985. Rearrangement of nitrogen fixation genes during heterocyst differentiation in the cyanobacterium *Anabaena*. Nature 314:419–423.

74. Forchhammer K, De Marsac NT. 1994. The P(II) protein in the cyanobacterium *Synechococcus* sp. strain PCC 7942 is modified by serine phosphorylation and signals the cellular N-status. J Bacteriol 176:84–91.

75. Timson DJ. 2005. Functional analysis of disease-causing mutations in human UDP-galactose 4-epimerase. FEBS J 272:6170–6177.

76. Beerens K, Soetaert W, Desmet T. 2013. Characterization and mutational analysis of the UDP-Glc (NAc) 4-epimerase from *Marinithermus hydrothermalis*. Appl Microbiol Biotechnol 97:7733–7740.

